# Gene expression comparisons between captive and wild shrew brains reveal captivity effects

**DOI:** 10.1101/2023.10.02.560583

**Authors:** Maria Alejandra Bedoya Duque, William R. Thomas, Dina K. N. Dechmann, John Nieland, Cecilia Baldoni, Dominik von Elverfeldt, Marion Muturi, Angelique P. Corthals, Liliana M. Dávalos

## Abstract

Compared to their free-ranging counterparts, wild animals in captivity experience different conditions with lasting physiological and behavioral effects. Although shifts in gene expression are expected to occur upstream of these phenotypes, we found no previous gene expression comparisons of captive vs. free-ranging mammals. We assessed gene expression profiles of three brain regions (cortex, olfactory bulb, and hippocampus) of wild shrews (*Sorex araneus*) compared to shrews kept in captivity for two months and undertook sample drop-out to examine robustness given limited sample sizes. Consistent with captivity effects, we found hundreds of differentially expressed genes in all three brain regions, 104 overlapping across all three, that enriched pathways associated with neurodegenerative disease, oxidative phosphorylation, and genes encoding ribosomal proteins. In the shrew, transcriptomic changes detected under captivity resemble responses in several human pathologies, including major depressive disorder and neurodegeneration. While interpretations of individual genes are tempered by small sample sizes, we propose captivity influences brain gene expression and function and can confound analyses of natural processes in wild individuals under captive conditions.

## INTRODUCTION

Wild animals placed in captivity are subject to altered environmental conditions in comparison to their free-ranging counterparts. Shelter, movement space, diet, climate, lighting, social factors, and human presence vary drastically between the wild and captivity [1–4]. Organisms respond to these changes by altering behavior, physiology, biochemistry and gene expression, but these responses occur at different time scales [5]. While long-term habituation to captivity can activate multiple stress responses, even short-term habituation to captive conditions affects brain morphology and function [1,3,6–8]. Symptoms of chronic stress appear because of frequent exposure to perceived threats [2] that with reduced environmental stimulation can lead to depressive-like states in captive animals [9]. Thus, despite the perception of a comfortable, predictable, and controlled environment, captive conditions can generate detrimental responses in captive individuals.

Short-term and long-term environmental changes also lead to lasting changes in gene expression, with a central role in cellular adaptation through extensive transcriptional and post-transcriptional regulation [10–13]. Although studies in mammals have addressed the effects of captivity in many phenotypes downstream of transcription, including morphology [14,15], longevity [16] and stress [17,18], to our knowledge, there are no experiments comparing brain gene expression in non-domesticated wild and captive mammals. This is of particular interest to modern research, as many experiments are performed on mammals in laboratory settings and may yield different results from those obtained in the wild [6,19], potentially confounding the effects of captivity with phenotypes of interest.

To explore potential changes in the brain from captivity, we assessed the brain gene expression profiles of wild common shrews (*Sorex araneus*) compared to same-age shrews held captive for two months (>10% of their lifespan). The common shrew is ideal for revealing potential captivity effects on gene expression, as many research resources are available: a complete genome assembly and a brain MRI-atlas with region-specific gene expression profiles [20]. *S. araneus* is also relevant to neuroscience for its seasonal change in brain size [21]. We focused on three brain regions that may shift functional importance in an environment that does not require exploring complex surroundings: 1) the cortex, whose evolution in mammals has improved information processing and decision making, 2) the hippocampus, involved in spatial memory and learning, and 3) the olfactory bulb, which receives olfactory information from the environment. We outline differences in brain gene expression between wild and captive shrews detected using differential gene expression analysis of mRNA. Our results support a captivity effect in the shrew brain, and we highlight similar responses found in studies of metabolism, neurodegenerative diseases, stress, and depression.

## MATERIALS AND METHODS

### Shrew Collection

Shrews were caught in Liggeringen, Germany, as previously described [22] to reduce stress. Our study had small sample sizes, as only ten individuals from the population could be sampled in autumn and logistical barriers from the COVID-19 pandemic restricted sampling over multiple years. Captive juvenile shrews were caught in September of 2020 and housed individually for two months at the Max Planck Institute of Animal Behavior before samples were taken (Table 1). Animals were allowed to move freely between two connected cages, one containing a running wheel, bowls of food and water, a nesting pot, and the other consisting of several layers of soil and hay (detailed housing in [23]). Captive shrews were kept at ambient outdoor temperatures, ranging from −5°C to 22°C, and provided a daily diet of 5g of raw meat and 2g of mealworm. Only four autumn wild juvenile shrews (two female and two male) were caught in November, and were sampled the same day of capture. Although sex is used as a covariate in subsequent analyses, there were two female wild individuals, but no captive females. This confounded sex and captivity in experimental design. Season was used as a proxy for age, as *S. araneus* have an unusually short lifespan of ∼13 months, with one reproductive season followed by senescence. Overlap between generations occurs in summer and early fall, when adults have worn teeth, disheveled fur, and developed gonads. Adults were excluded from the experiment.

**Table 1.**
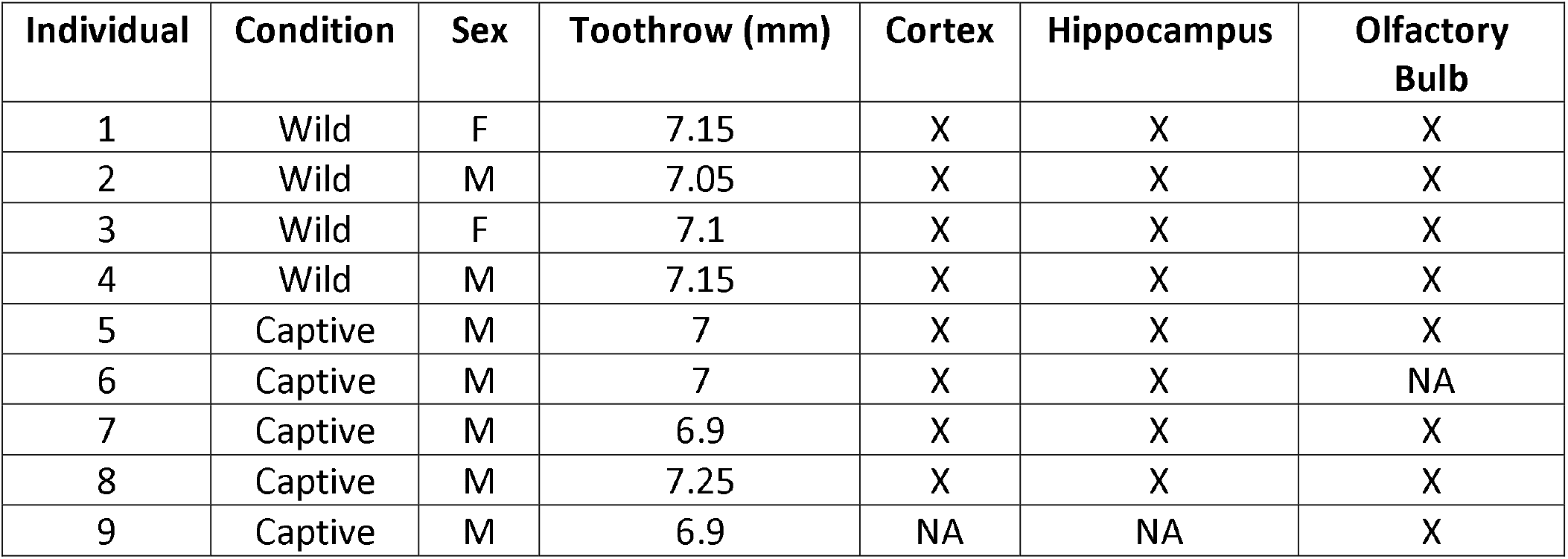
Descriptions of individuals used in gene expression experiments.

### Tissue Sampling, RNA Extraction, and Sequencing

Shrews were weighed, then anesthetized and euthanized under deep anesthesia by perfusion of the vascular system with PAXgene Tissue Fixative (perfusion described in Lázaro et al., 2018 [22]). The brain was extracted immediately, weighed, dissected into individual regions, and submerged in cold fixative. Three brain regions were compared between captive and wild individuals: cortex, olfactory bulb, and hippocampus. Brain regions were held in fixative, and within 24 hours were switched to PAXgene Tissue Stabilizer and frozen in liquid nitrogen. Skull toothrow length was measured to assess the size of individuals. RNA was extracted from each brain region using a modified Qiagen Micro RNAeasy protocol designed to retrieve RNA from small sensory tissues [24]. Quality control, library preparation, and sequencing were conducted by Genewiz. RNA integrity numbers (RIN) were measured with RNA ScreenTape and samples below a RIN of 6 were removed from the experiment. mRNAs were selected by polyA enrichment followed by paired-end sequencing with Illumina HiSeq.

### Bioinformatics

Samples with low sequencing depths (<20 million reads) were removed from the experiment. Adapters were trimmed and low-quality reads removed with fastp 0.20.1 [25]. Reads were mapped by pseudo- alignment to the *S. araneus* reference transcriptome (GCF_000181275.1_SorAra2.0_rna) using Kallisto 0.46.1 [26]. Samples with mapping percentage under 45% were removed. Read counts were then normalized for library size and content using the median of ratios from DESeq2 1.36.0 [27]. To visualize the overall effect, we ran a principal component analysis (PCA) of the 500 most varying genes for each brain region using variance stabilized read counts [28].

Differential gene expression between captive and wild individuals for each brain region was analyzed in R (version 4.2.0) with DESeq2 [27]. Sex and body size were used as additional covariates for the analysis (∼sex+toothrow+condition). Maxillary tooth row length was used as a proxy for body size, as it does not change seasonally within individuals [29,30]. Statistical significance for each region comparison was determined by conducting a modified Wald test, using an alternative null of 20% change, and then adjusting p-values for multiple testing using Benjamini and Hochberg correction [31] (p_adj_ < 0.05). The proportion of true nulls was analyzed with limma [32]. Given a statistical significance threshold of 0.05, the expected proportion of true nulls approaches 0.95. Higher values indicate false negatives, and lower values indicate false positives. Genes identified as differentially expressed for each brain region were separated by direction (upregulated or downregulated). KEGG pathway enrichment analysis (Modified Fisher’s Exact test) was carried out for both directional gene lists of each brain region using DAVID [33,34]. To examine result robustness, we conducted a drop-out analysis for each brain region (Supplemental Figures 1-3). We removed one individual per analysis, identified overlapping differentially expressed genes across all analyses, reran DAVID pathway enrichments, and identified overlapping pathways.

**Figure 1.**
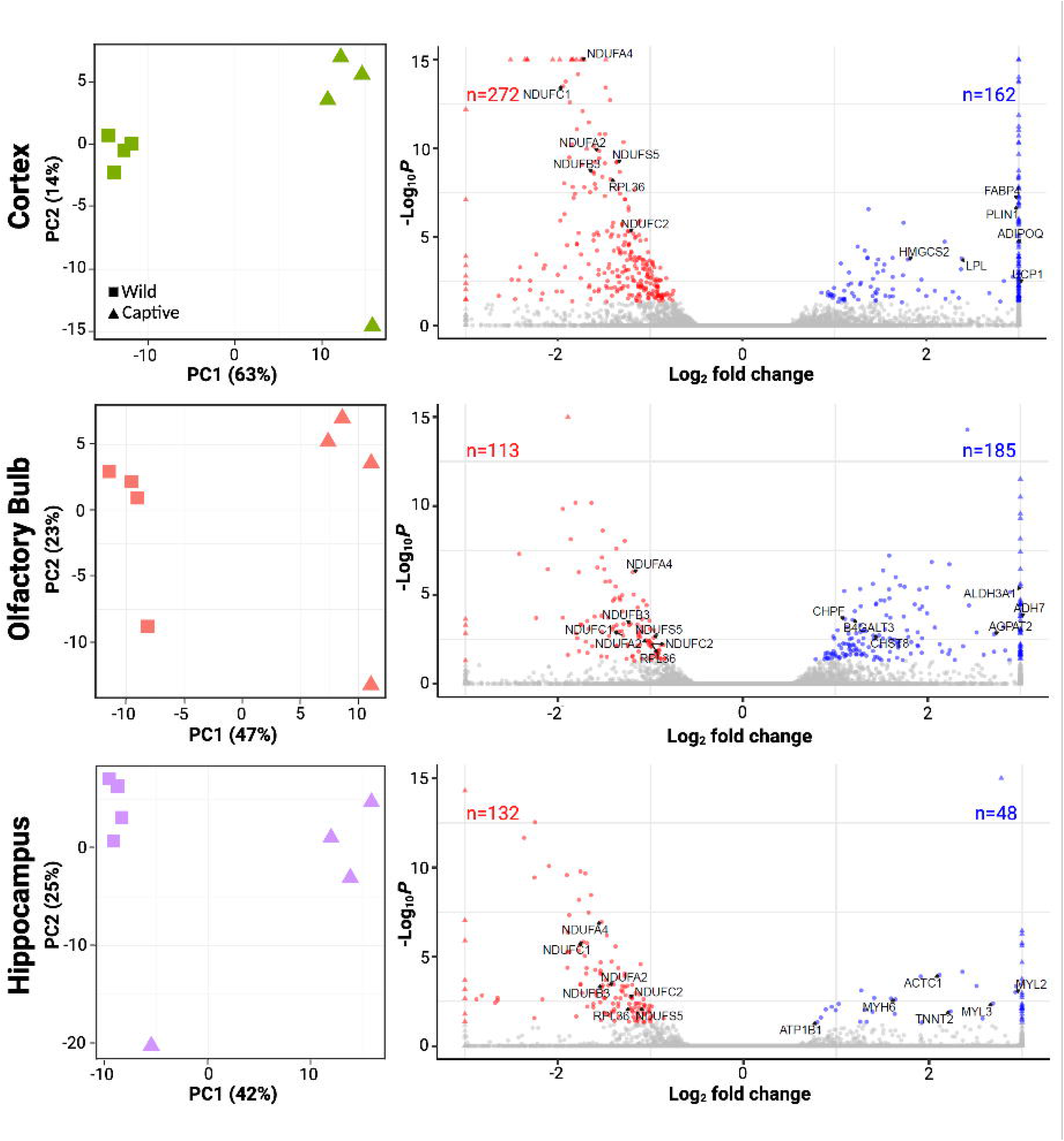
Differential gene expression between wild (n=4) and captive (n=4) shrews in the cortex, olfactory bulb, and hippocampus. *Left:* PCA of the top 500 varying genes for each brain region. *Right*: Volcano plots for the cortex, olfactory bulb, and hippocampus gene expression. Significant (p_adj_ < 0.05) DEGs are shown in red (downregulated) or blue (upregulated) relative to wild, DEGs of interest are labeled. Labeled downregulated genes encode for ribosomal proteins and mitochondrial complex I subunit proteins, which have also been differentially expressed in previous studies. Labeled upregulated genes correspond to the top 6 genes from the most significant upregulated enriched pathways.

## RESULTS

We identified divergence in the expression profiles of all three brain regions associated with a potential captivity effect. For each brain region, PCA produced two principal component axes that accounted for most of the variation from these genes (Figure 1). The first principal component separated wild from captive individuals, accounting for 42% of the variation in gene expression in the hippocampus and 47% in the olfactory bulb. Percent variation expressed was even greater in the cortex, where the first principal component accounted for 63% of the variation. As for PC2, each region highlighted one sample within each analysis with variation not found in other samples. We determined this was not a single outlier sample, as we found each case corresponded to different individuals for each brain region (Supplemental Figure 4), with no discernible pattern across regions.

Differential expression analysis confirmed these patterns (Figure 1). Of the 19,296 genes analyzed, 603 genes were differentially expressed (DEGs; p_adj_<0.05) in at least one brain region (Figure 2). In the cortex, we found 434 DEGs, 162 upregulated and 272 downregulated in captive shrews. The olfactory bulb had 298 DEGs, with 185 upregulated and 112 downregulated in captive animals. In the hippocampus, we identified 180 DEGs, 48 upregulated and 132 downregulated in captive animals. The proportion of true nulls was high, 0.97 in the cortex, 1.00 in the hippocampus, and 0.99 in the olfactory bulb. This indicates lower statistical power to detect many true positives in the hippocampus and olfactory bulb because of small samples sizes and high variability [35,36]. We compared lists of DEGs between brain regions and found 104 genes that were differentially expressed in the same direction across all regions, with another 101 overlapping in two of three brain regions (Figure 2).

**Figure 2.**
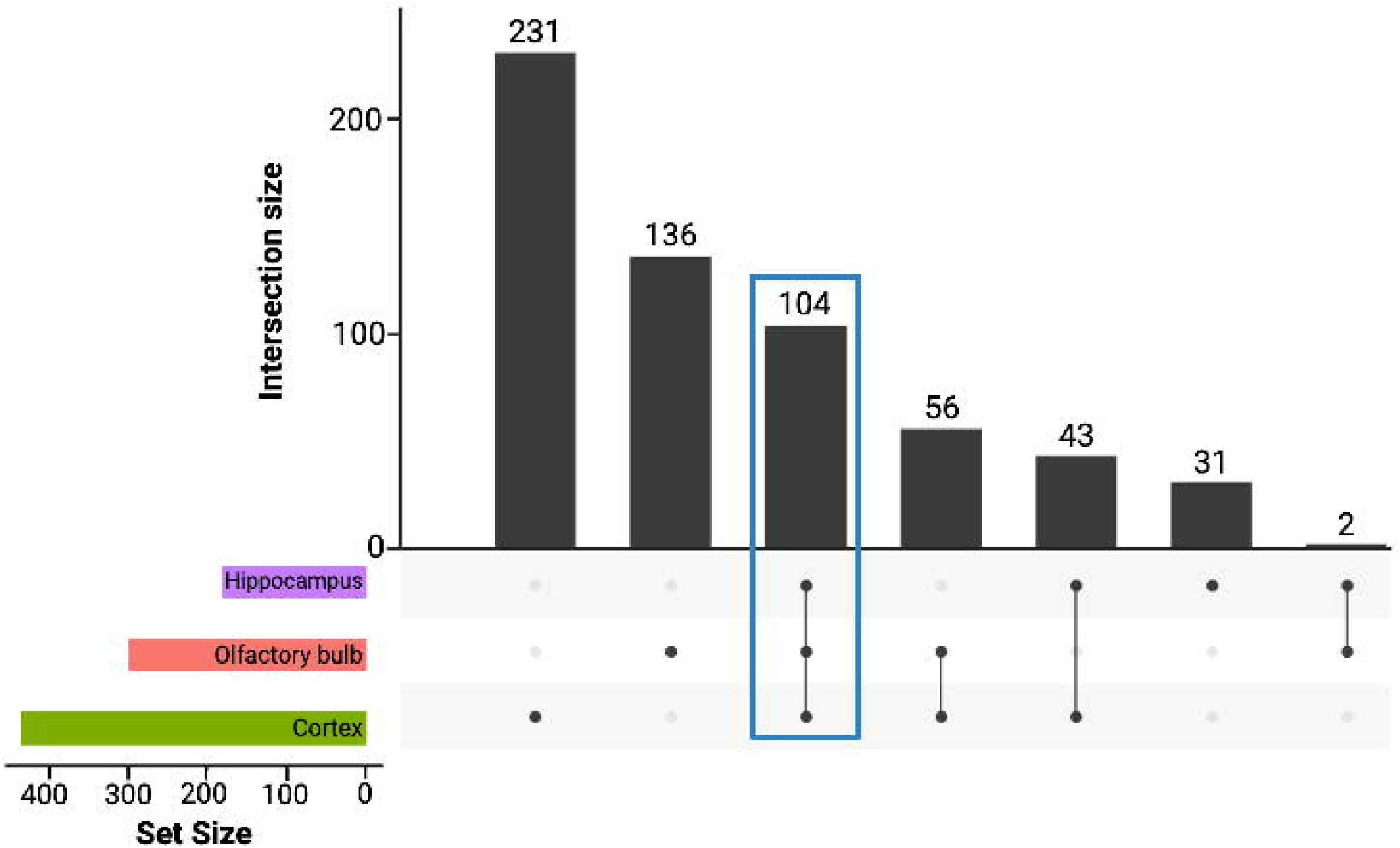
Overlapping DEGs for the three brain regions. The side histogram corresponds to the number of DEGs for each brain region (set size), and the top histogram corresponds to intersections between the sets of DEGs across the three brain regions. All the 104 DEGs that overlap between the three brain regions (highlighted in blue) have the same directional change in gene expression (79 downregulated, 25 upregulated).

To partially overcome small sample sizes, we undertook pathway enrichment analyses, which revealed captive gene expression differences are linked to ribosomal and mitochondrial function, with associations to neurodegenerative processes (Figure 3). We identified 21 pathways significantly enriched (p-value < 0.05, modified Fisher’s Exact Test) for the cortex, 21 for the olfactory bulb and 21 for the hippocampus. Out of these, 14 pathways were significantly enriched (p<0.05) across the three brain regions (Figure 3), with the largest enrichment in the ribosome pathway (p _cor_: 7.4 × 10^−32^, n= 35genes; p _olf_: 4.0 × 10^−11^, n=13 genes; p _hip_ : 5.0 × 10^−15^, n=17 genes). Within enriched pathways, we identified expression changes that likely affect mitochondrial function in the brain. Nineteen genes encoding mitochondrial complex I NADH:ubiquinone oxidoreductase subunit proteins (largest enzyme of the respiratory chain), of which 8 overlap all three regions and were found across 12 of the 14 overlapping enriched pathways (Figure 3). Six of the pathways containing complex I proteins were associated with neurodegenerative diseases, including Parkinson’s disease, amyotrophic lateral sclerosis, pathways of neurodegeneration-multiple diseases, prion disease, Huntington’s disease, and Alzheimer’s disease. Out of the eight enriched pathways remaining, six of them were associated with changes in metabolism, such as oxidative phosphorylation, retrograde endocannabinoid signaling, non-alcoholic fatty liver disease, diabetic cardiomyopathy, chemical carcinogenesis - ROS, and thermogenesis, highlighting the influence of mitochondrial genes on both metabolically related and neurodegenerative mechanisms.

**Figure 3.**
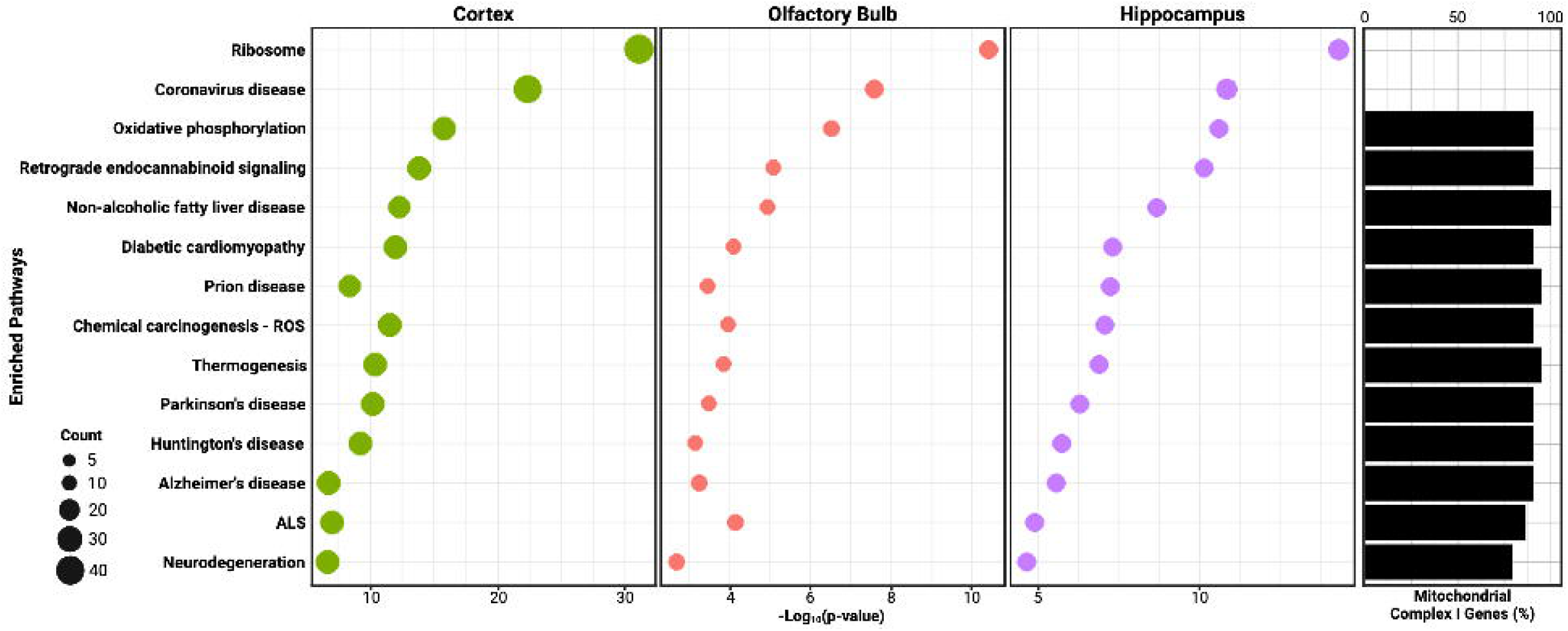
*Left:* Pathway analysis for captive and wild shrews for the cortex, olfactory bulb and hippocampus. These pathways were significantly enriched across the three brain regions when analyzing downregulated genes, and were also enriched when analyzing only the 104 overlapping significant genes. *Right:* Side histogram represents percentage of complex I subunit genes in each of these pathways.

While principal component analyses show little impact from sex, drop-out analyses indicate robust gene expression results in the cortex (Supplemental Figures 5 and 1-3). First, we examined the expression of each individual mitochondrial gene of interest for each brain region (Supplemental Figures 6A-B). Normalized counts did show some dependance on sex. But instead of driving differences in gene expression, sex differences appear to dampen overall signal, and differences are greater between wild and captive samples than between sexes. Pathway enrichment for drop-out analyses was robust for the cortex, where 109 significant genes from the full experimental analysis remained differentially expressed across all dropouts (p_adj_<0.05). This subset of robust genes enriches most of the proposed pathways, including oxidative phosphorylation, neurodegeneration, Parkinson’s disease, Alzheimer’s disease, prion disease, Huntington’s disease, and ALS (p<0.05). The olfactory bulb and hippocampus were not robust to sample drop-out, as only 27 genes for the hippocampus and 15 genes for the olfactory bulb remained across all drop-out analyses and did not enrich highlighted pathways.

## DISCUSSION

Both genomic [37,38] and environmental variation [39–41] impact neural structures and function. Past studies have focused on fixed domestication-associated neural changes in subsets of mammals [42–44], and identified differences in gene expression and transcription factors involved in domestication-associated changes in both brain and craniofacial phenotypes [37,38]. While brain gene expression responses to captive environment change have been documented in diverse organisms [45–48], this is the first study documenting potential captivity effects specifically on mammalian brain transcripts in the absence of artificial selection. Although captivity is routine in many experiments, effects on expression may parallel or counter responses to domestication or other forms of induced or natural environmental change. Potentially large expression changes in the brain after a brief stint in captivity highlight the importance of working with both wild and captive individuals to avoid confounding gene expression changes from phenotypes of interest with those associated with captivity.

We found potential region-specific responses to captivity, but small sample sizes mean these results are limited by low statistical power and should be interpreted with caution. Principal components separate samples by captivity first (Figure 1) so that overall expression supports a captivity effect. Individual DEGs, however, are not robust to sample drop-out for the underpowered hippocampus and olfactory bulb (Supplemental Figure 2 and 3). For example, in the hippocampus we found apparent upregulation of four myosin genes (*MYH6, MYL2, MYL3, MYBPC3*) (Figure 1). This may relate to changes in synaptic plasticity, as myosins regulate actin cytoskeleton dynamics in dendrites essential for long-term potentiation and memory formation in rodents [49–52]. But based on power analyses from vertebrate brain expression [36], when sampling four individuals, there is ∼70% power to accurately detect differential expression in genes with more than 40 reads. While most observed DEGs are above this threshold (Supplemental Figure 7), nearly half of the total genes sequenced had fewer than 40 reads, reducing the power to detect differential expression (∼5% to 45%), and therefore the reliability for each individual differentially expressed gene.

Instead, we focus biological interpretations on pathway changes found across all three brain regions (Figure 3), as gene sets reduce noise and are more stable compared to individual gene expression changes [53]. We found downregulation of ribosomal protein genes in captive shrews. Ribosomes compose the cell’s protein synthesis machinery, and their own synthesis, ribosomal biogenesis, is a highly conserved process [54,55]. In the captive shrew brain, downregulation of 12 ribosomal proteins (RPs) suggests impaired regulation of ribosomal biogenesis and subsequent functions. By determining translational capacity of proliferating cells [56], ribosomal biogenesis constrains animal growth and development [57,58]. Dysregulation of ribosomal biogenesis is therefore implicated in numerous phenotypes including neurite survival and synaptic function associated with human neurodegeneration [54]. These effects seem to be mirrored in captive shrews, as we found downregulated gene enrichment for pathways associated with neurodegeneration. Neurodegeneration has been linked to changes in stress [59,60], but these transcriptomic profiles and potential shifts in processes such as reduced synaptic efficiency might be occurring because of captivity and not disease.

Differential expression of ribosomal protein-coding genes in the brains of captive shrews could influence behavior, as many such genes have been previously highlighted for their regulatory changes under domestication and social defeat stress [44,61]. For example, comparisons between domestic rabbits and wild counterparts identified 612 DEGs across four brain regions that also enriched ribosomal pathways [44]. Additionally, changes in expression of ribosomal genes have also been associated with decreased proliferation in the hippocampus of mice under chronic social defeat stress, which consists of repeated negative social experiences (defeats) produced by daily agonistic interactions with an aggressive partner, leading to states of anxiety and depression [61]. Captivity-associated differential expression of ribosomal proteins in the shrew brain may thus indicate a depressed neural state, especially as past studies have argued that animals in captivity often experience repeated stressors used to induce depressive-like states [9].

Captivity conditions such as altered diet and physical activity may also influence shrew energy metabolism in the brain. In captive shrews, we found downregulation of mitochondrial complex I proteins, essential for energy production, as they comprise the first and largest complex of the electron transport chain [62,63]. Impairments to complex I function are linked to pathophysiological states, genetic diseases [62,64–66], and several neurodegenerative disorders, leading to decline of energy production and increased production of reactive oxygen species [67–70]. Downregulation of 8 genes encoding for mitochondrial complex I subunit proteins in three shrew brain regions suggests a dysfunction of the oxidative phosphorylation (OXPHOS) system and dysregulation of energy metabolism in captivity. These genes are also downregulated in the brains of patients with autism [71], and neurodegenerative diseases such as Huntington’s [72] and Alzheimer’s [73]. Indeed, the effects of certain conditions found in captivity, such as reduced stimulation, could also modify OXPHOS function in association with neurodegeneration, as normal brains exposed to chronic sensory deprivation exhibit downregulation of OXPHOS subunits as found in the brain of Alzheimer’s patients [74]. However, studies have found differences in mitochondrial function/gene expression between sexes [75–77], and sex- dependent susceptibility to neurodegenerative disease [76]. Therefore, including only two female samples is a limitation of our study to be improved in the future.

As with differential ribosomal protein expression, mitochondrial dysfunction and changes in expression of complex I subunits have also been linked to depressed activity and stress-related behaviors. For example, female mice bred for low activity showed upregulation of complex I subunit genes [78], including three genes downregulated in the three brain regions tested in captive shrews (*NDUFA2, NDUFA4*, and *NDUFS5*). These genes exhibit opposite expression patterns between low activity mice and captive shrews. But other studies have found the direction of changes in mitochondrial gene expression related to stress and anxiety may depend on stressor duration, genetic background, and strain of the model organism studied [79]. Nevertheless, published and newly detected shifts still point to mitochondrial pathway dysregulation as a major brain stress response [79]. Differential expression of complex I subunit genes in captive shrews is consistent with a transcriptional phenotype associated with depression and stress in model organisms. While the validation of specific mechanisms for such a transcriptional phenotype (e.g., by western blot validation) is outside the scope of this paper, we have identified key potential targets for future analyses.

## Conclusion

Despite attempts to limit differential stressors and optimize holding conditions, these results indicate transcriptomic changes in the brain of shrews after being placed in a captive environment. Such changes could influence phenotype through shifts in energy metabolism, synaptic function, cell survival, and behavior. Across pathways, captivity-associated changes in the shrew resemble responses in several human diseases, notably, neurodegeneration. These are unlikely to be shrew-specific, as similar expression shifts have been found in other organisms with disturbance in their natural environments. When investigating the molecular mechanisms of disease using model organisms, captivity can confound analyses of natural physiological processes. There are benefits to using captive organisms in disease research, but our results indicate future studies should use natural environments to illuminate brain function and responses to environmental change. These experiments could involve collaborative efforts with ecologists and evolutionary biologists [80] to study natural models of brain change [21,81] that can be validated in laboratory settings using cell lines. Alternatively, findings from laboratory-based disease models can be paired with experimental groups in exclosures as close to wild as possible to assess the impact of environmental factors on experimental outcomes.

## Supporting information

Electronic Supplementary Materials

## ACKNOWLEDGMENTS

Images were created with a licensed version of BioRender.com.

## SUPPLEMENTAL MATERIAL

Supplemental Figs. S1–S7, supplemental table 2: https://doi.org/10.6084/m9.figshare.27237300.v1

Supplemental Table 1: DOI: https://doi.org/10.5061/dryad.qz612jmng

