## Supplementary material for "Gene expression comparisons between captive and wild shrew brains reveal captivity effects": Electronic Supplementary Materials: SupplementalFigure1.pdf

### Cortex

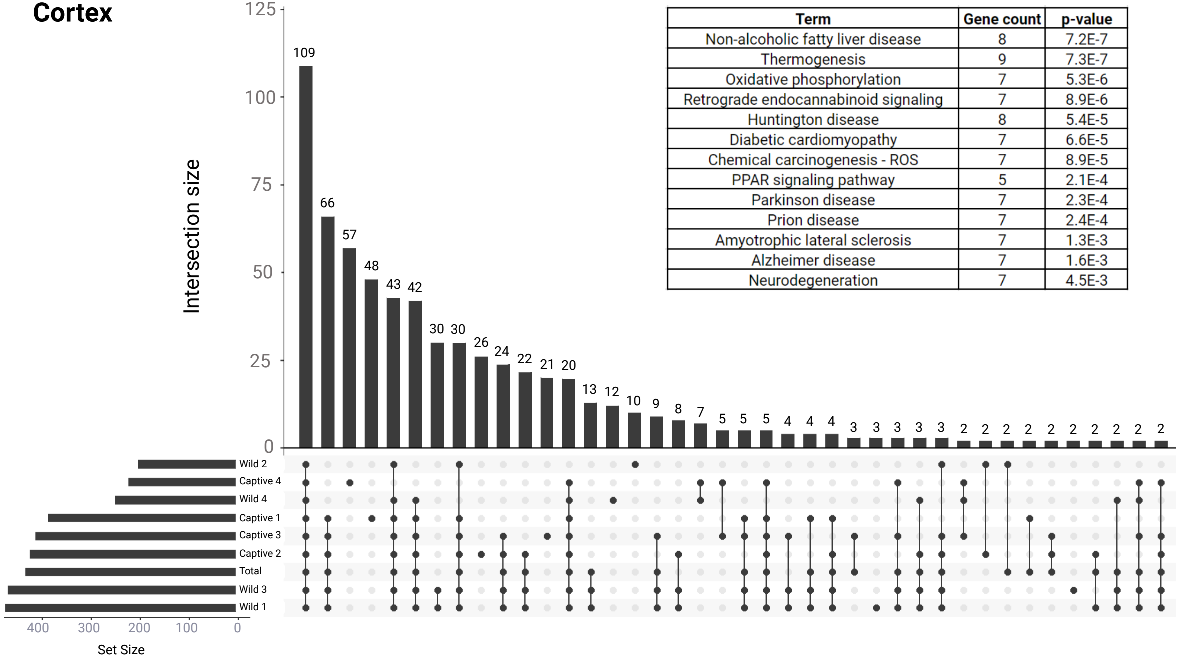


| Term | Gene count | p-value |
|---|---|---|
| Non-alcoholic fatty liver disease | 8 | 7.2E-7 |
| Thermogenesis | 9 | 7.3E-7 |
| Oxidative phosphorylation | 7 | 5.3E-6 |
| Retrograde endocannabinoid signaling | 7 | 8.9E-6 |
| Huntington disease | 8 | 5.4E-5 |
| Diabetic cardiomyopathy | 7 | 6.6E-5 |
| Chemical carcinogenesis - ROS | 7 | 8.9E-5 |
| PPAR signaling pathway | 5 | 2.1E-4 |
| Parkinson disease | 7 | 2.3E-4 |
| Prion disease | 7 | 2.4E-4 |
| Amyotrophic lateral sclerosis | 7 | 1.3E-3 |
| Alzheimer disease | 7 | 1.6E-3 |
| Neurodegeneration | 7 | 4.5E-3 |
