## Supplementary material for "Gene expression comparisons between captive and wild shrew brains reveal captivity effects": Electronic Supplementary Materials: SupplementalMaterial.docx

**Supplemental Figure 1.**

Leave-one-out analysis for the cortex testing 19,296 genes (n=3 to 4 captive vs n=3 to 4 wild individuals). Side histogram corresponds to number of DEGs after dropping one sample for each analysis (set size), and the top histogram corresponds to intersections between sets of DEGs in each analysis. 109 genes were robust to excluding samples. Table corresponds to pathway enrichment for robust genes for each analysis. Discussed pathways of interest such as Oxidative phosphorylation, Parkinson’s disease, Prion disease, Alzheimer’s disease, Huntington’s disease, Amyotrophic lateral sclerosis (ALS), and Pathways of neurodegeneration - multiple diseases (Neurodegeneration) are still enriched after leave-one-out analysis.

**Supplemental Figure 2.**

Leave-one-out analysis for the olfactory bulb testing 19,296 genes (n=3 to 4 captive vs n=3 to 4 wild individuals). Side histogram corresponds to number of DEGs after dropping one sample for each analysis (set size), and the top histogram corresponds to intersections between sets of DEGs in each analysis. 15 genes were robust to excluding samples. Pathways discussed for the olfactory bulb were not robust to the drop-out analysis, suggesting reduced effect of captivity for this brain region.

**Supplemental Figure 3.**

Leave-one-out analysis for the hippocampus testing 19,296 genes (n=3-4 captive vs n=3-4 wild individuals). Side histogram corresponds to number of DEGs after dropping one sample for each analysis (set size), and the top histogram corresponds to intersections between sets of DEGs in each analysis. Only 27 genes were robust to excluding samples, which did not enrich any of the notable pathways.

**Supplemental Figure 4.**

*Top*: Principal component analysis of the top 500 varying genes between wild (n=4) and captive (n=4) shrews in the cortex, olfactory bulb, and hippocampus, colored by individual samples. Different samples deviate along PC2 for each brain region. *Bottom:* Loading plot of the principal component analysis for each brain region. No discernible pattern is observed for PC2 through the three brain regions.

**Supplemental Figure 5.**

Principal component analysis of the top 500 varying genes between wild (n=4) and captive (n=4) shrews in the cortex, olfactory bulb, and hippocampus, colored by sex. Axes show the percentage of the variance each component accounts for. Although sex is confounded with captivity, there is minimal impact of sex in the wild set compared to captive effect.

**Supplemental Figure 6A-B.**

Plots of normalized counts for the eight differentially expressed complex I subunit protein genes found across 12 of the 14 overlapping enriched pathways for the three brain regions. Y axis corresponds to gene expression (normalized counts) and X axis represents the division between captive and wild samples. Female samples (n=2) are colored blue, male samples (n=6) are colored yellow. Although mitochondrial gene expression is sex-dependent, greater difference in gene expression can be observed between wild and captive samples, than between male and female samples.

**Supplemental Figure 7.**

Distribution of reads per gene (n=19,296) for all brain regions analyzed. Significant DEGs are filled with blue; 268 in the olfactory bulb, 382 in the cortex, and 161 in the hippocampus. Most significant DEGs have high (~70%) power. As some genes reside under the 40-read threshold, increased sample size can improve their results.

**Supplemental Table 2-** Pathway enrichment for the 104 DEGs for all 3 regions

| **Term** | **Gene Count** | **PValue** | **Genes** | **Fold**  **Enrichment** | **Bonferroni** | **Benjamini** | **FDR** |
| --- | --- | --- | --- | --- | --- | --- | --- |
| Ribosome | 12 | 8,27E-10 | RPS15A, RPS29, RPL34, RPLP1, RPL36, RPL35A, RPL38, RPL27, RPL37, RPS21, RPL39, RPL7 | 12,57315 | 3,97E-08 | 3,97E-08 | 3,14E-08 |
| Coronavirus disease - COVID-19 | 12 | 9,45E-09 | RPS15A, RPS29, RPL34, RPLP1, RPL36, RPL35A, RPL38, RPL27, RPL37, RPS21, RPL39, RPL7 | 9,975706 | 4,54E-07 | 2,27E-07 | 1,8E-07 |
| Oxidative phosphorylation | 8 | 4,52E-07 | NDUFA4, NDUFS5, NDUFA3, NDUFA2, NDUFB3, NDUFC2, NDUFC1, ATP6V1F | 15,7536 | 2,17E-05 | 7,23E-06 | 5,73E-06 |
| Retrograde endocannabinoid signaling | 7 | 1,29E-05 | NDUFA4, NDUFS5, NDUFA3, NDUFA2, NDUFB3, NDUFC2, NDUFC1 | 12,70327 | 0,000619 | 0,000155 | 0,000123 |
| Non-alcoholic fatty liver disease | 7 | 1,79E-05 | NDUFA4, NDUFS5, NDUFA3, NDUFA2, NDUFB3, NDUFC2, NDUFC1 | 11,99753 | 0,000858 | 0,000172 | 0,000136 |
| Diabetic cardiomyopathy | 7 | 9,22E-05 | NDUFA4, NDUFS5, NDUFA3, NDUFA2, NDUFB3, NDUFC2, NDUFC1 | 8,956682 | 0,004417 | 0,000738 | 0,000584 |
| Chemical carcinogenesis - reactive oxygen species | 7 | 0,000124 | NDUFA4, NDUFS5, NDUFA3, NDUFA2, NDUFB3, NDUFC2, NDUFC1 | 8,487336 | 0,005938 | 0,000851 | 0,000674 |
| Thermogenesis | 7 | 0,000153 | NDUFA4, NDUFS5, NDUFA3, NDUFA2, NDUFB3, NDUFC2, NDUFC1 | 8,166387 | 0,00733 | 0,00092 | 0,000728 |
| Prion disease | 7 | 0,000323 | NDUFA4, NDUFS5, NDUFA3, NDUFA2, NDUFB3, NDUFC2, NDUFC1 | 7,119414 | 0,015366 | 0,001579 | 0,00125 |
| Parkinson disease | 7 | 0,000329 | NDUFA4, NDUFS5, NDUFA3, NDUFA2, NDUFB3, NDUFC2, NDUFC1 | 7,093431 | 0,015668 | 0,001579 | 0,00125 |
| Huntington disease | 7 | 0,000613 | NDUFA4, NDUFS5, NDUFA3, NDUFA2, NDUFB3, NDUFC2, NDUFC1 | 6,31039 | 0,02902 | 0,002676 | 0,002119 |
| Amyotrophic lateral sclerosis | 7 | 0,001705 | NDUFA4, NDUFS5, NDUFA3, NDUFA2, NDUFB3, NDUFC2, NDUFC1 | 5,182933 | 0,078629 | 0,006819 | 0,005398 |
| Alzheimer disease | 7 | 0,002054 | NDUFA4, NDUFS5, NDUFA3, NDUFA2, NDUFB3, NDUFC2, NDUFC1 | 4,996401 | 0,09397 | 0,007583 | 0,006003 |
| Pathways of neurodegeneration - multiple diseases | 7 | 0,005872 | NDUFA4, NDUFS5, NDUFA3, NDUFA2, NDUFB3, NDUFC2, NDUFC1 | 4,040748 | 0,246258 | 0,020134 | 0,015939 |
