## Supplementary figures and images for "Gene expression comparisons between captive and wild shrew brains reveal captivity effects"

### SupplementalFigure2.pdf

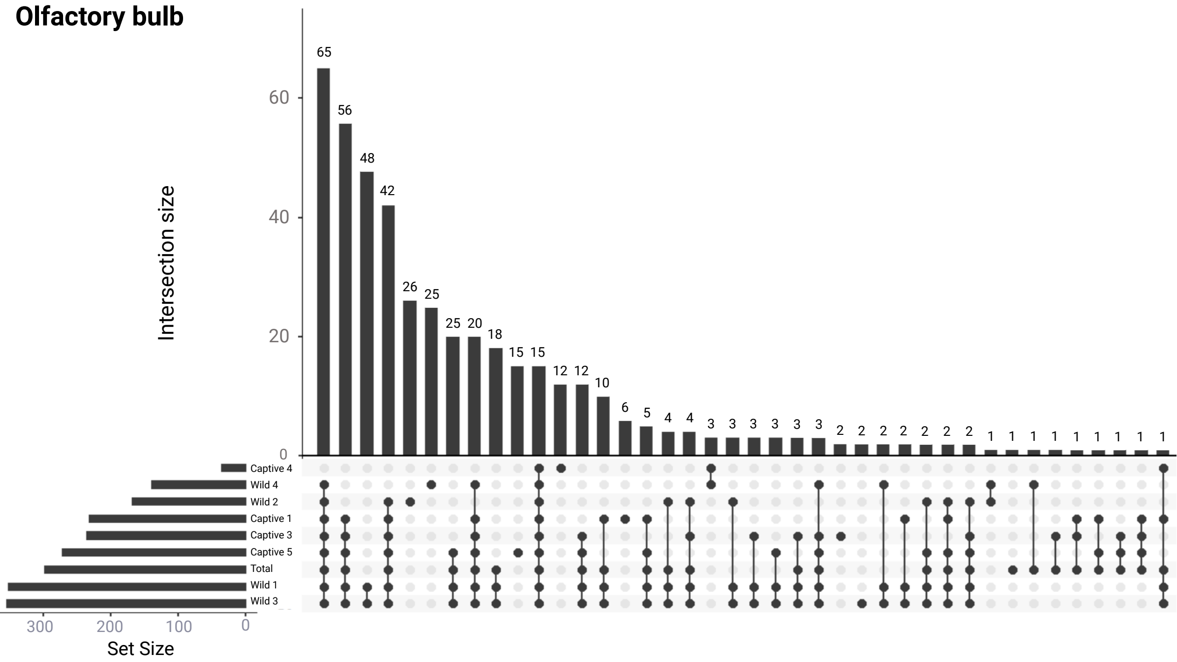

### SupplementalFigure3.pdf

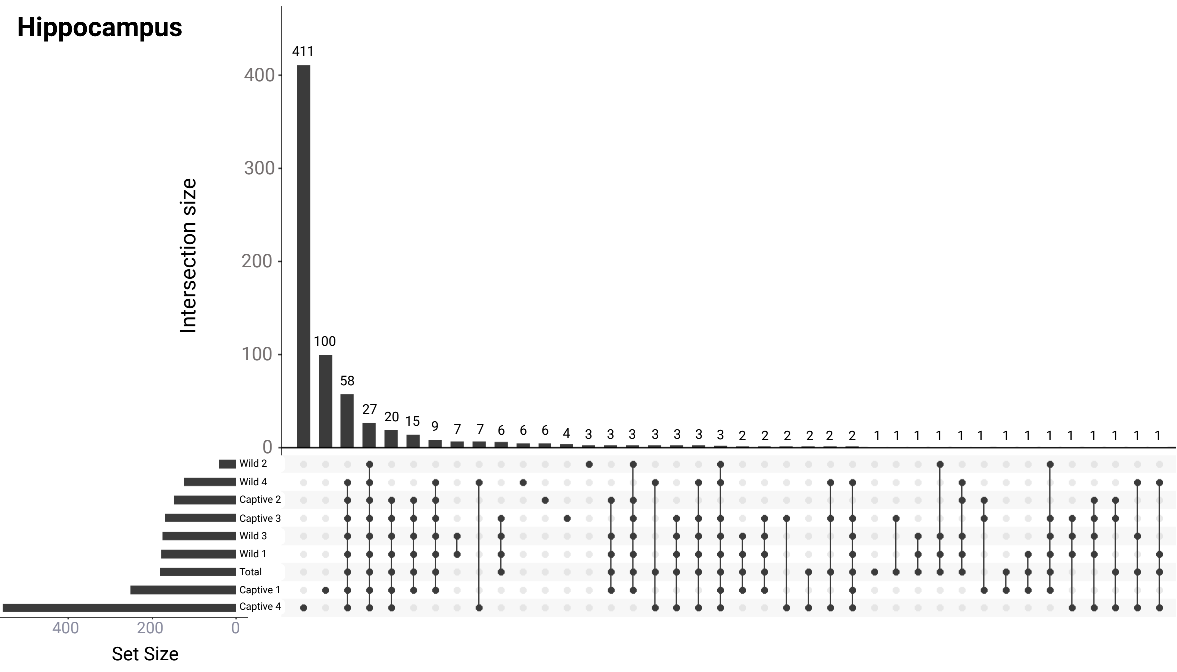

### SupplementalFigure4.pdf

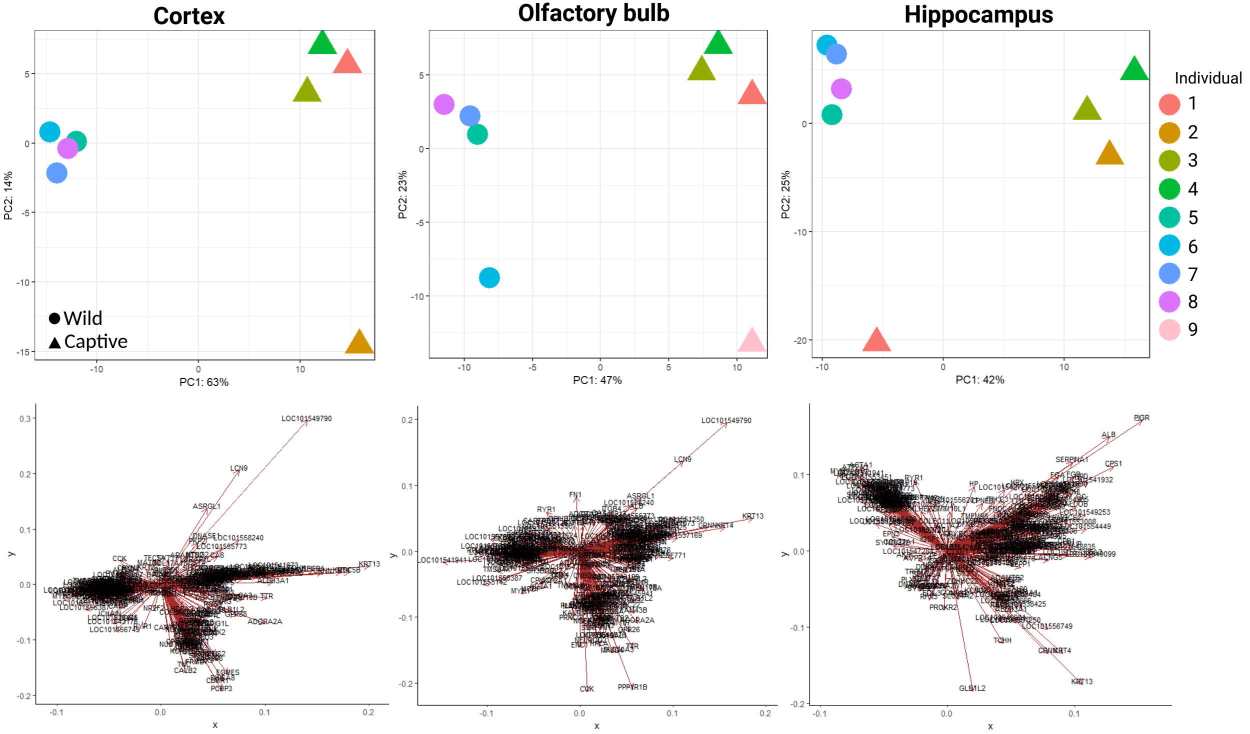

### SupplementalFigure5.pdf

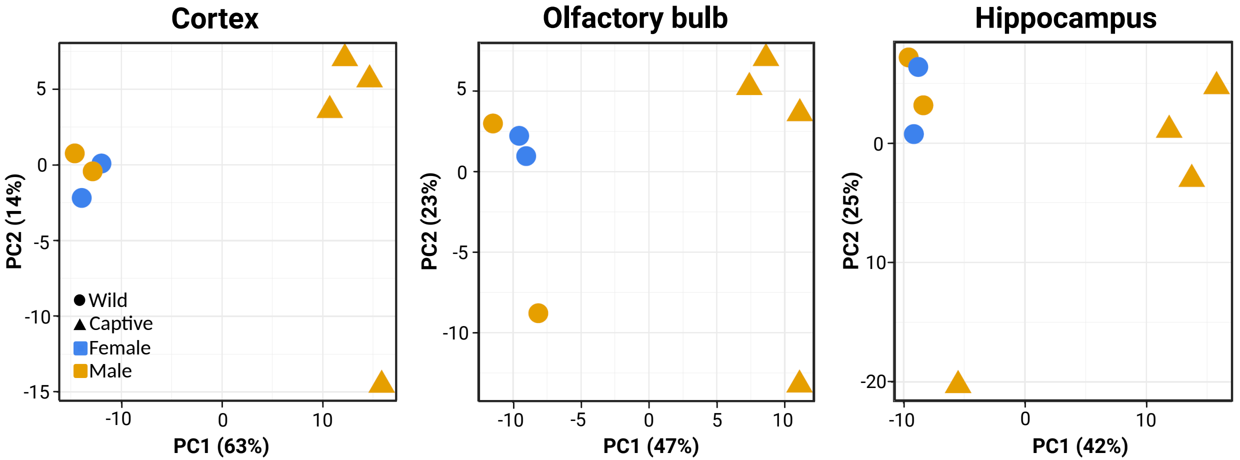

### SupplementalFigure6A.pdf

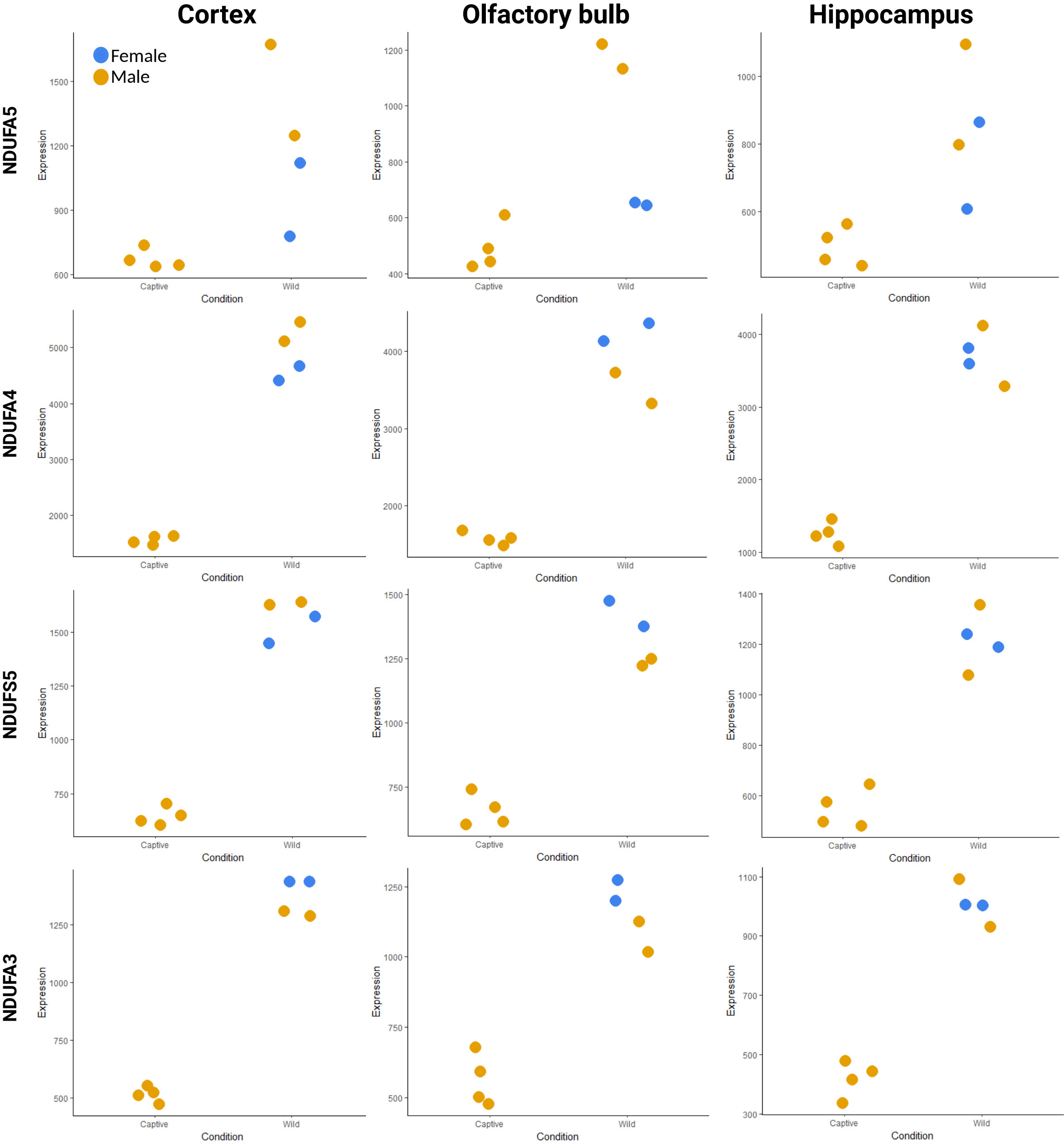

### SupplementalFigure6B.pdf

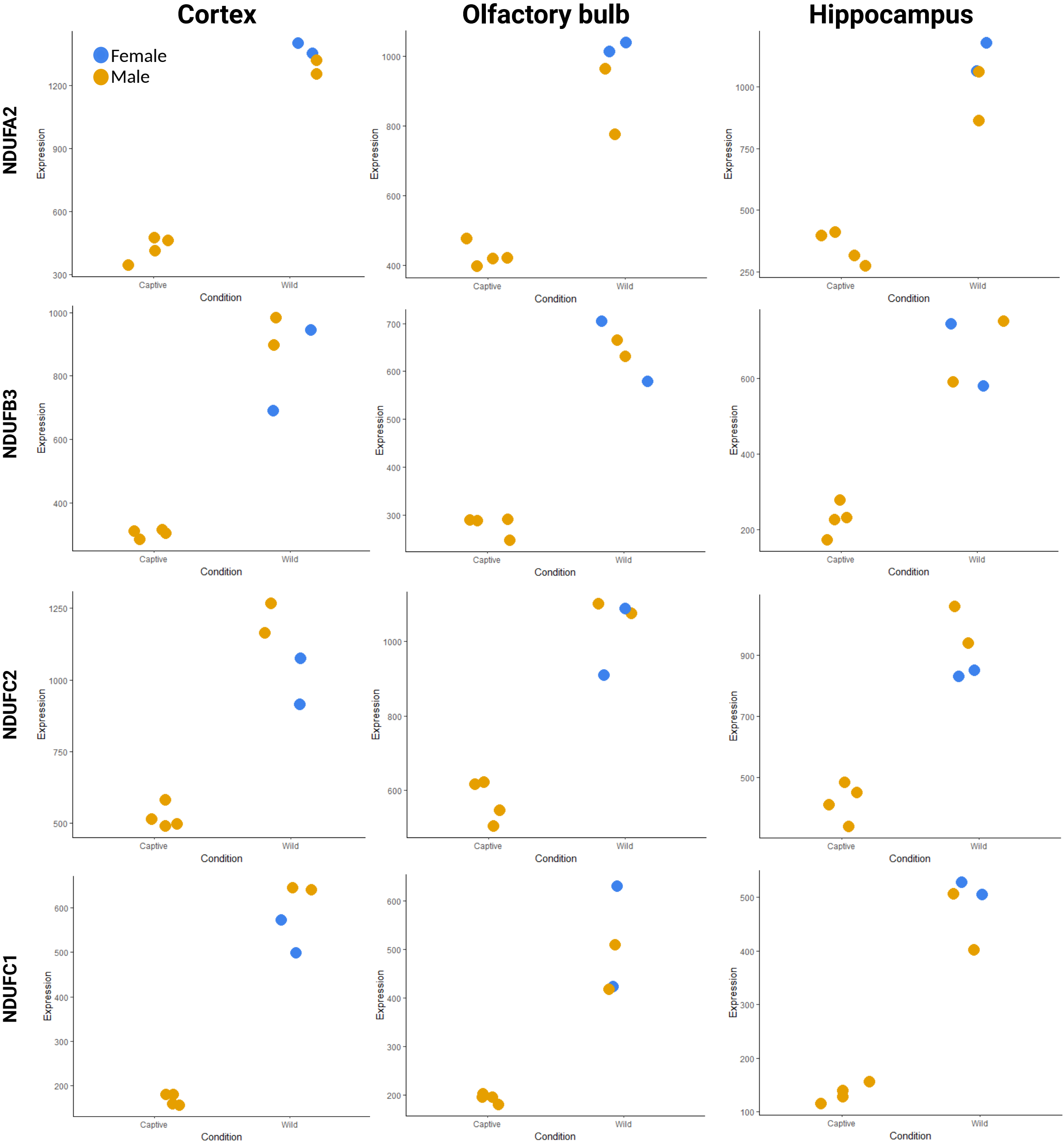

### SupplementalFigure7.pdf

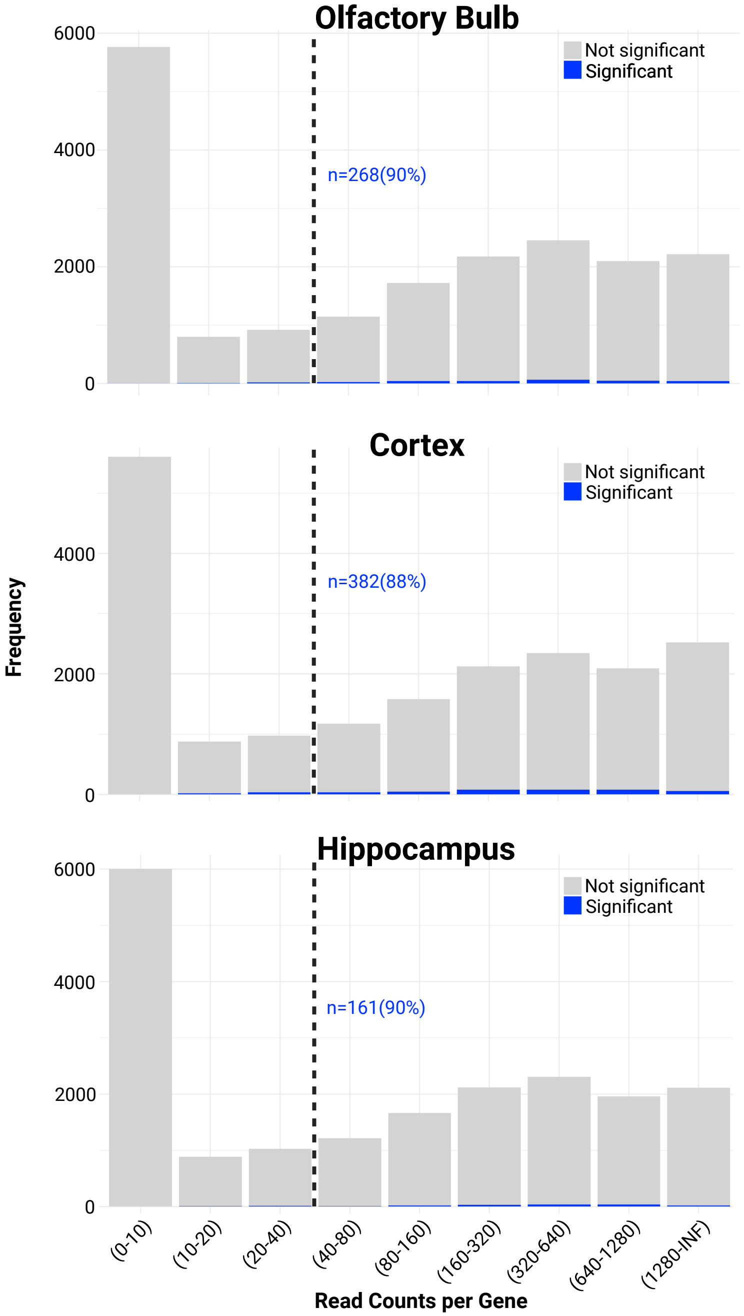
